# Climate change drives contrasting redistribution patterns in endemic and endangered Himalayan *Gentiana*

**DOI:** 10.64898/2026.06.03.729779

**Authors:** Syed Waseem Gillani, Mushtaq Ahmad, Muhammad Manzoor, Raja Waqar Ahmed Khan, Amir Sohail, Roberto Salguero-Gómez

## Abstract

1. Rapid climate warming threatens mountain biodiversity, particularly species with narrow climatic niches and limited dispersal capacity. Mountain ecosystems are especially vulnerable because steep environmental gradients restrict opportunities for species redistribution. The Kashmir Himalaya, a globally important biodiversity hotspot experiencing accelerated warming, has already undergone an increase of approximately 0.8 °C during the 20th century and is projected to warm by 2.5–2.8 °C by the 2050s. Despite climatic changes, future persistence of many threatened plants remains poorly understood.
2. Here, we evaluate present and future habitat suitability for two Himalayan taxa, *Gentiana cachemirica*, an endemic species, and *Gentiana kurroo*, a critically endangered species. Specifically, we quantify how climate change and topographic variability influence species distribution and persistence under multiple emission scenarios.
3. We applied species distribution models (SDMs) to presence data of both species and forecasted habitat suitability under four Shared Socioeconomic Pathways (SSP126, SSP245, SSP370, and SSP585). We hypothesized that *G. cachemirica*, a narrow-niche, low-dispersal species, is expected to lose habitat due to thermal sensitivity, while *G. kurroo* may persist or expand under favorable scenarios because of broader tolerance and higher dispersal. We also expected microclimatic refuges to buffer populations, whereas extreme warming would accelerate habitat decline.
4. The predictions of our SDMs under current conditions indicate a highly restricted and fragmented habitat for *G. cachemirica*, covering 651 km², but a broader suitable area (2,452 km²) for *G. kurroo*. In agreement with our hypotheses, our forecasts indicate severe habitat contraction (55–70%) for *G. cachemirica* across all SSPs, but scenario-dependent responses for *G. kurroo*, including modest expansion under low-emission scenarios and severe declines under high-emission scenarios. Centroid analyses suggest pronounced climate-driven range shifts, with *G. kurroo* projected to migrate up to 33 km toward the east-southeast by 2100, while *G. cachemirica* is projected to display limited dispersal capacity.
5. **S*ynthesis*.** Our findings suggest that climatic niche breadth, dispersal limitation, and topographic buffering strongly mediate species responses to warming in mountain ecosystems. Endemic specialists are projected to experience disproportionate habitat fragmentation and range restriction, highlighting the importance of conserving climatic refugia and elevational connectivity under rapid environmental change.

## 1. Introduction

The on-going accelerating climate change is driving a rapid decline in global biodiversity (Bellard et al., 2012; Pimm, 2014; Ishaq et al., 2016; Kattel, 2022; Dainese et al., 2024). Indeed, current extinction rates far exceed background levels, putting up to one million species at risk and highlighting the need to identify the main drivers of biodiversity loss (Didham et al., 2007; Barnosky et al., 2011; Pimm et al., 2014; IPBES, 2019; Ceballos et al., 2017). Multiple interacting pressures are contributing to these declines, and differences in their roles across regions often limit the effectiveness of conservation planning (Bellard et al., 2016; Maxwell et al., 2016; Howard et al., 2020). Mountain ecosystems are especially sensitive, as warming increases thermal stress and ecological vulnerability at higher elevations (Rangwala & Miller, 2012; Pepin et al., 2015; Stoffel & Huggel, 2012). Rising temperatures also shift species distributions and reshape communities, further increasing extinction risk (Pecl et al., 2017; IPCC, 2021; Rubenstein et al., 2023; Dainese et al., 2024; Khan et al., 2025; Gillani et al., 2026). These changes highlight the urgent need for reliable ecological predictions to guide effective conservation in a rapidly changing world (Mouquet et al., 2015; Khan et al., 2025).

Species distribution models (SDMs), also known as ecological niche or habitat suitability models, are fundamental tools in ecology, biogeography, and conservation science (Elith & Leathwick 2009; Franklin 2010; Guisan et al. 2017; Peterson et al. 2011). These models integrate species occurrence data with environmental variables to generate spatially explicit predictions of habitat suitability across space and time (Marsh et al., 2023). SDMs enable reconstruction of historical distributions and forecasting range dynamics under climate change (Guisan & Thuiller 2005; Maiorano et al. 2013; Gould et al., 2014; Yang et al., 2023). Anticipating biodiversity responses to climate change represents a central objective in contemporary ecological research (Reyer et al., 2013; Akhmedov et al., 2021). Climate-based simulations enhance predictive accuracy and reveal key environmental drivers shaping distributional change (Zeng et al., 2022; Bernatchez et al., 2023). SDMs therefore provide essential guidance for conservation planning, restoration prioritization, and management of threatened species under accelerating environmental change (Guillera-Arroita et al., 2015; Fois et al., 2016; Sanz-Arnal et al., 2022).

The Himalayas are a globally important mountainous biodiversity hotspot that is facing rapid and severe climate change impacts, threatening its ecological stability (An, 2001; Zhu et al., 2025; Gillani et al., 2025). The accelerated warming in the Himalayas compared to the global average has intensified cryosphere loss, glacier retreat, biodiversity decline, and has also reduced its essential ecosystem services (Shrestha et al., 2012; Lamsal et al., 2017; Rani et al., 2022). Glacier decline represents a dominant regional threat, with serious consequences for downstream water availability and long-term livelihood security across Asia (Bolch et al., 2012; Immerzeel et al., 2012; You & Xu 2022). Moreover, intensified human pressures, including deforestation, habitat fragmentation, agriculture, pollution, and resource overuse, are further increasing the ecosystem vulnerability and degradation in the region (Aryal & Kerkhoff 2008; Gopirajan et al., 2022). Surprisingly, despite supporting millions of people, the Himalayan region remains critically data-deficient, limiting robust assessment of species-level impacts and conservation planning (IPCC, 2007; Gautam et al., 2013; Sabin et al., 2020).

Rare and endangered plants are highly sensitive to environmental change. This high sensitivity is partly due to their low abundance, restricted range, and specialized habitat requirements (Markham, 2014; Rejmánek, 2018). These constraints limit their ability to tolerate climatic variation and reduce their capacity to disperse to new suitable areas (Broennimann et al., 2006; Casazza et al., 2014). Several studies show that ecological specialization can push species toward extinction when environmental conditions change (Preston et al., 2008; Williams et al., 2009; Işik, 2011). This pattern is well documented in range-restricted rare and endemic plants that decline rapidly under climate change (Casazza et al., 2014; Sękiewicz et al., 2024; Gillani et al., 2026). Therefore, understanding the current and potential geographic distributions of range-restricted species is important for conservation planning, as this knowledge is key to identify vulnerable populations and anticipate climate-driven range shifts (Guisan et al., 2013; Khan et al., 2025; Manzoor et al., 2026).

Here, we assess how climate change may affect the habitats of two gentian species of differing conservation status (*Gentiana cachemirica* is endemic, *G. kurroo* is critically endangered). To do so, we use SDMs with climatic and topographic predictors. We specifically test four hypotheses: (H1) Rising maximum temperature of the warmest month (BIO5) and lower precipitation of the driest month (BIO14) will shrink and fragment *G. cachemirica* populations, confining them to high-elevation, north-facing refugia due to its narrow niche, thermal sensitivity, and limited dispersal. Similar contractions of cold-adapted mountain plants under warming have been widely reported (Kelly & Goulden, 2008; Gentili et al., 2015a; Rumpf et al., 2018). (H2) *G. kurroo* will persist or even expand under low-emission scenarios by tracking suitable climates to higher elevations and east-southeast directions. This response is driven by its broader climatic tolerance and border niche breadth. Species with broader ecological niches generally show greater capacity to track shifting climatic envelopes and maintain viable populations under moderate climate change (Thuiller et al., 2005; Slatyer et al., 2013). However, under high emissions, rising maximum temperature of the warmest month (BIO5), increased temperature annual range (BIO7), precipitation seasonality (BIO15), steep slopes, and south-facing aspects will reduce habitat suitability and fragment populations of *G. kurro* due to warmer and drier microclimates. (H3) Comparing the two taxa, *G. cachemirica* will show minimal range shifts and greater habitat loss due to its narrow niche, whereas *G. kurroo* will exhibit scenario-dependent expansions toward higher elevations and east-southeast directions, driven by broader niche breadth and higher dispersal capacity. These differences reflect the niche breadth-range size relationship, where species with broader climatic tolerances respond more effectively to environmental change (Brown, 1995; Slatyer et al., 2013; Evans & Jacquemyn, 2022). Finally, (H4) areas with moderate temperatures, high dry-season precipitation, north-facing slopes, and gentle topography will act as stable refugia for both species. In contrast, extreme warming, high temperature variability, steep slopes, and south-facing aspects will cause habitat loss and fragmentation. These expectations are consistent with ecological theory linking restricted ranges and niche specialization to higher climate sensitivity (Swihart et al., 2003; Duarte et al., 2019; Fekete et al., 2023).

## 2 Methodology

To test our hypotheses on climate-driven distribution changes, we modelled the current and future habitat suitability of two Himalayan species, *Gentiana cachemirica* and *G. kurroo*. We combined species occurrence data with key climatic and topographic variables to predict potential distributions under multiple climate scenarios. We then quantified habitat shifts, centroid movements, and fragmentation to compare species responses and evaluate the roles of niche breadth, dispersal capacity, and topographic refugia.

### 2.1 Study area

The study area is located in northern Pakistan (31° 30′ - 37° 00′ N, 69° 00′ - 77° 30′ E; Fig. 1). Climatic, topographic, and edaphic factors shape vegetation patterns in this region. Indeed, elevation and temperature exert strong control over plant distribution (Bahuguna et al., 2016). Rainfall shows marked spatial variation across the region, with the Eastern areas receiving summer monsoon rainfall from June to September, but the Northern and western zones mainly depend on winter precipitation from December to March (Khan et al., 2025). The summer monsoon contributes approximately 60% of annual rainfall (Khan et al., 2025). Most areas experience arid to semiarid conditions, with less than 250 mm of rainfall per year (Khan et al., 2025). In contrast, northern mountainous areas receive between 760 and 2,000 mm annually. High mountains and extensive glaciers produce severe winter conditions, with temperatures occasionally reaching −50 °C (Khan et al., 2025).

**Figure 1.**
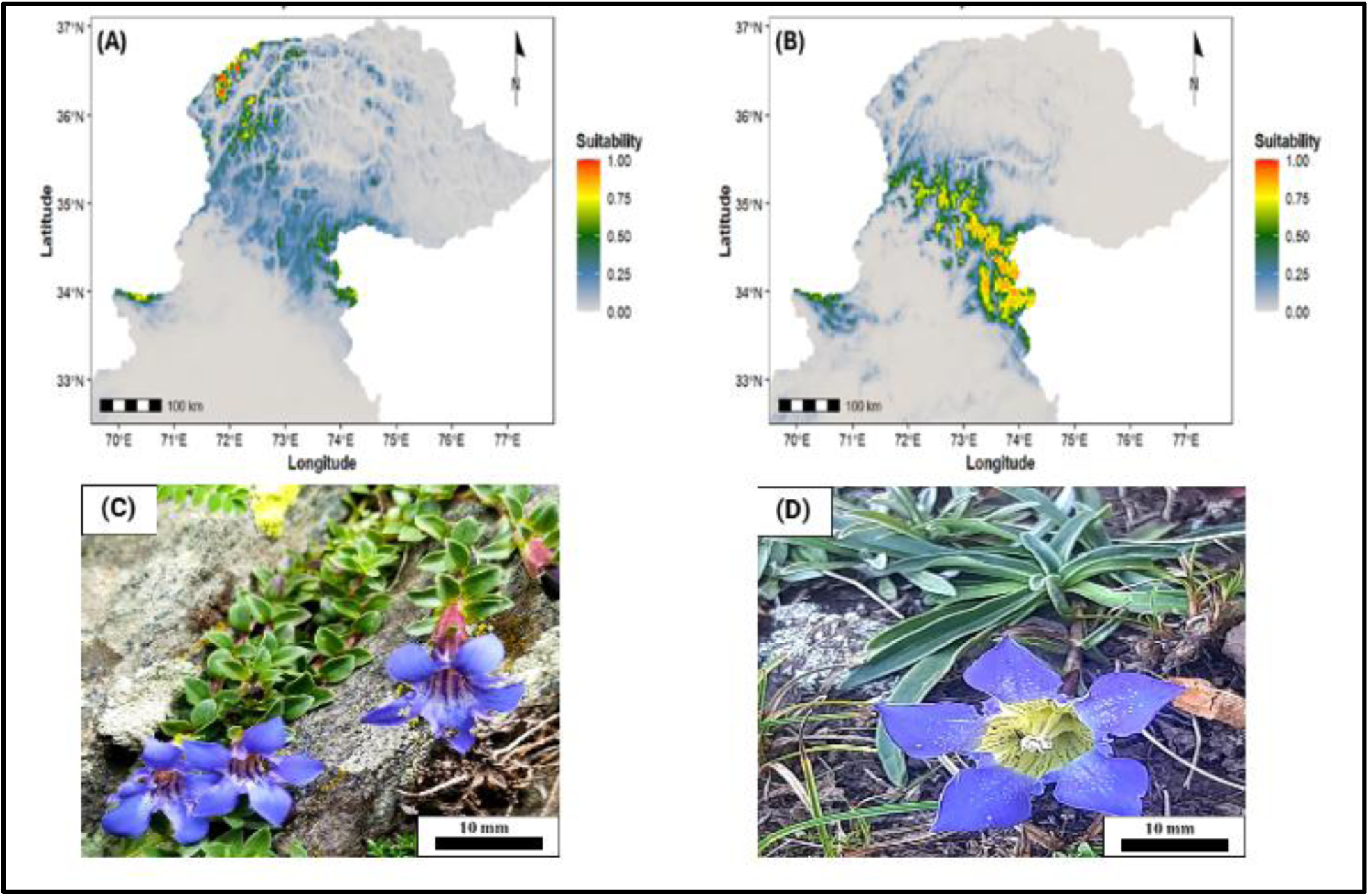
Baseline habitat suitability patterns reveal clear differences in distribution and niche breadth between *Gentiana cachemirica* and *G. kurroo* in the Western Himalayas. (A) Predicted current habitat suitability for *G. cachemirica* under baseline climatic conditions (1970–2000) shows fragmented and spatially restricted suitable habitat concentrated in high-elevation regions. (B) Predicted current habitat suitability for *G. kurroo* shows a broader and more continuous distribution across valley systems and mid-elevation zones. Habitat suitability values range from 0 (unsuitable; gray) to 1 (highly suitable; red), derived from generalized linear models incorporating climatic and topographic predictors. (C) Field photograph of *G. cachemirica* growing in rocky alpine habitat under natural conditions. (D) Field photograph of *G. kurroo* showing characteristic growth form and flowering morphology.

### 2.2 Study system

Genus *Gentiana* (Gentianaceae) includes approximately 400 annual and perennial herb species distributed across Europe, Asia, and northern Australia (Editorial Committee of Flora of China 1988; Luo et al., 2026). Species of *Gentiana* typically inhabit alpine screes, meadows, and shrublands (Zhengyi 1987). *Gentiana* plants contain abundant secondary metabolites, including iridoids, secoiridoids, and related derivatives, which support their medicinal importance. Gentiopicroside and swertiamarin are the principal bioactive compounds, providing traditional therapeutic functions such as heat clearing and regulation of hepatic and biliary activity (Commission 2025). A gap in comparative species distribution modelling for endemic *Gentiana cashmirica* and critically endangered *Gentiana kurroo* positions this research as the first assessment from Pakistan.

We focus on two Himalayan species of the genus Gentiana, *Gentiana cachemirica* and *G. kurroo*, which occupy distinct ecological environments. *Gentiana cachemirica*, an endemic species of the Kashmir Western Himalayas, grows in alpine habitats characterized by heavy snowfall, low temperatures, and strong seasonal variability. Field observations show that the species occurs mostly above 3,300 m a.s.l., where it establishes small populations within sun-exposed rock crevices. In comparison, *G. kurroo* inhabits temperate zones and grows on open, sunny slopes under relatively moderate climatic conditions. The species flowers mainly from July to October.

### 2.3 Species occurrence data and spatial alignment

To test our hypotheses, we generate high-quality inputs for distribution modelling, involving the curation and data filtering of occurrence records of *Gentiana cachemirica* and *G. kurroo*. Specifically, we obtained occurrence records from the Global Biodiversity Information Facility (GBIF; https://www.gbif.org/) liturature and field surveys in the Kashmir Himalaya. To reduce the effects of spatial autocorrelation and sampling-effort bias, we used the spThin package (Aiello-Lammens et al., 2015) in R (v. 4.5.2; R Core Team, 2025). To mitigate spatial autocorrelation, account for geographic sampling bias, and maintain consistency with the spatial resolution of the environmental predictors, we applied a minimum nearest-neighbor distance of 1 km among occurrence records. This filtering process yielded a curated dataset of 30 and 48 high-quality records for *G. cachemirica* and *G. kurroo*, respectively. We standardized all coordinates to the WGS84 geographic datum to standardize the spatial reference across all data sources and prevent coordinate misalignment during environmental data extraction. To ensure model integrity, we performed a final extraction check. We retained occurrences overlapping valid cells only to ensure model consistency and to avoid issues with missing predictor data (Mainali et al., 2020; Hijmans & Elith, 2023).

### 2.4 Environmental predictors

To ensure accurate, bias-minimized inputs for distribution modelling, we assembled and refined species occurrence records. The initial environmental data comprised 22 predictors, including 19 bioclimatic variables and five topographic variables. We obtained the baseline climatic data (representing the 1970-2000 period) from the WorldClim database (Fick & Hijmans, 2017). Next, we sourced the bioclimatic variables (BIO1-BIO19) directly from WorldClim (https://www.worldclim.org/). Last, we derived topographic predictors including elevation (altitude above sea level), slope (the steepness of the terrain), and aspect (the compass direction a slope faces) from elevation data to capture terrain-related influences on the distribution of our two target species.

We modelled the future habitat suitability of *G. cachemirica* and *G. kurroo* using climate projections, topographic layers, and harmonized environmental covariates. To model future habitat suitability, we used projections from the ACCESS-CM2 general circulation model (Bi et al., 2020) within the Coupled Model Intercomparison Project Phase 6 framework. This modelling follows scenarios reported by the Intergovernmental Panel on Climate Change (IPCC, 2021). These projections accounted for four Shared Socioeconomic Pathways (SSP126, SSP245, SSP370, and SSP585) across three temporal slots: 2041-2060, 2061-2080, and 2081-2100. We developed pertinent topographic layers using WorldClim digital elevation data as a primary template, with subsequent layers being computed via the *terra* package in R (Hijmans et al., 2025). To ensure computational consistency, we harmonized all environmental covariates to a 30 arc-second spatial resolution and clipped to the specific geographical extent of the study area.

### 2.5 Species presence and pseudo-absence data

We prepared presence-only data with optimized background points and filtered environmental variables for accurate species modelling. The modelling relied on presence-only data, so we randomly sampled background points across the study area (Barbet-Massin et al., 2012; Phillips et al., 2009), treating these as pseudo-absence data. Ecological pseudo-absences generation improves species distribution modelling (Broussin et al., 2024). Random pseudo-absence (background) sampling is a widely used and effective baseline strategy in species distribution modelling, often demonstrating strong performance (Descombes et al., 2022; Broussin et al., 2024).To determine an appropriate background sample size, we iteratively tested various background sizes for each species, ultimately selecting the densities that maximized individual model performance. Once the spatial coordinates were established, we extracted environmental values for all presence and pseudo-absence localities. We cleaned the resulting dataset by removing records with incomplete predictor data or missing raster values to ensure analytical consistency, following established protocols (Barbet-Massin et al., 2012; Senay et al., 2013)

### 2.6 Predictor screening and collinearity

To minimize collinearity among environmental variables and ensure reliable parameter estimation, we filtered predictors for zero variance and high correlation, standardizing remaining variables to select parsimonious sets for each species. To minimize redundancy and mitigate multicollinearity, we subjected the initial predictor pool to a two-stage filtration protocol. First, we identified and subsequently removed variables with zero variance using the *caret* R package (Kuhn, 2014). We then standardized all remaining predictors using a z-score transformation, centering the data based on the mean and standard deviation of each variable. In the second stage, we assessed correlations among environmental variables nby calculating pairwise Pearson correlation coefficients (Pearson, 1895; Benesty et al., 2009), considering variables with a correlation coefficient greater than 0.7 as highly correlated. For each correlated pair, we removed one variable to reduce redundancy while retaining the most informative predictor. Addressing multicollinearity is critical in regression-based species distribution models (SDMs), as it stabilizes coefficient estimates and ensures that spatial inferences remain statistically sound (Dormann et al., 2013). As such, this refinement process resulted in a parsimonious final set of predictors, comprising four variables (BIO5, BIO14, Slope, and Aspect) for *G. cachemirica* and five variables (BIO5, BIO7, BIO15, Slope, and Aspect) for *G. kurroo* used in the modelling.

### 2.7 Species distribution modelling and assessment

To model species-environment relationships, we fitted separate generalized linear models (GLMs) for each species. To test our hypotheses (H1 and H2), where we predicted that increasing temperature and reduced precipitation would constrain and fragment *G. cachemirica* to cooler, high elevation refugia (H1), and whether *G. kurroo* would show relative persistence or range shifts under moderate conditions but declining suitability and fragmentation under more extreme warming (H2). Accordingly, we modelled the relationships between environmental predictors and species occurrence for *G. cachemirica* and *G. kurroo* separately.

To do so, we first transformed predictors into probabilities via a logit link and calibrating against multiple pseudo-absence datasets (Barbet-Massin et al., 2012; Guisan et al., 2017; Hijmans et al., 2024). We separately fitted SDMs for *G. cachemirica* and *G. kurroo* using GLM in the R package *dismo* (McCullagh, 2019; Hijmans et al., 2024). We modelled species occurrence using a set of environmental predictors combined linearly through regression coefficients (McCullagh & Nelder, 1989; Elith & Leathwick, 2009; Cho et al., 2022). We then back-transformed the resulting values via a logit link to estimate the probability of species presence.

We quantified the predictive accuracy of our models using multiple complementary metrics, including the Area Under the Receiver Operating Characteristic Curve (AUC; Hanley & McNeil, 1982), sensitivity and specificity (Fleiss et al., 2013), and the True Skill Statistic (TSS; Allouche et al., 2006). We used the AUC to assess the model’s overall ability to discriminate between presence and absence locations, where values range from 0.5 (no better than random) to 1 (perfect discrimination), and values above 0.7 generally indicate acceptable performance. We calculated sensitivity (true positive rate) and specificity (true negative rate) to measure the proportions of correctly predicted presences and absences, respectively; both metrics range from 0 to 1, with values closer to 1 indicating higher predictive accuracy. We used the TSS as a threshold-dependent metric to balance omission and commission errors, as it is insensitive to prevalence. We calculated TSS as sensitivity + specificity − 1, with values ranging from −1 to +1, where values ≤ 0 indicate no predictive skill and values above 0.5 are typically considered indicative of good model performance. We determined the optimal probability threshold for binary classification by maximizing Youden’s J statistic (Youden, 1950), defined as J = sensitivity + specificity − 1. J ranges from 0 to 1, with higher values indicating better combined performance of sensitivity and specificity, and its maximum identifies the threshold that optimally separates presences from absences.

We estimated how each environmental factor influences species distributions and model performance. To do so, we quantified the importance of each variable on each species’s distribution using standardized regression coefficients (Schielzeth, 2010) and then its permutation importance (Altmann et al., 2010), which evaluates the decrease in model performance following randomization of predictor values. We used standardized coefficients to assess the relative strength and direction of predictor effects within the fitted models. Then, we applied permutation importance to measure the contribution of each predictor to overall model performance. We calculated permutation importance as the decrease in the AUC after randomly permuting the values of a given predictor variable, thereby disrupting its relationship with species occurrence (Bring, 1994; Thuiller et al., 2009; Bladen & Cutler, 2024). Larger reductions in AUC following permutation indicated greater importance of the corresponding predictor for model discrimination ability.

### 2.8 Habitat suitability projection and spatial dynamics

We projected current and future habitat suitability to quantify changes in distribution, area, and potential range shifts. We calibrated the final models using the full standardized dataset and implemented them in the *dismo* R package (Elith & Leathwick, 2009; Hijmans et al., 2024). We projected habitat suitability under current and future climatic conditions, resulting in continuous probability maps ranging from 0 to 1. Next, we derived binary habitat suitability maps by applying the probability threshold that maximized Youden’s J statistic (Youden, 1950). We estimated the total suitable area of each species by multiplying the number of suitable raster cells by the area of an individual pixel. Next, we categorized changes in habitat suitability between current and future projections such as habitat gain, loss, or stability based on transitions in binary suitability status.

To test our hypothesis H3, where we predicted that *G. cachemirica* would exhibit limited range shifts and greater habitat loss, whereas *G. kurroo* would show comparatively greater spatial redistribution and potential range expansion toward higher elevations and east-southeast directions under changing climatic conditions, we compared projected changes in habitat suitability, range extent, and spatial distribution between the two species. To assess spatial shifts in species distributions, we calculated centroids of highly suitable habitat and used this information to quantify the direction and magnitude of projected range shifts (Loarie et al., 2009). Finally, we computed distances and bearings between centroids using great-circle geometry to characterize the magnitude and direction of potential migration. Briefly, great-circle geometry computes the shortest path between two points on the Earth’s curved surface using geographic coordinates, thus providing accurate estimates of distance and direction (bearing) between locations (Snyder, 1987; Bullock, 2007). We conducted all spatial analyses and visualizations in R using raster-based workflows (Hijmans et al., 2025), ggplot2 (Wickham, 2016), dplyr (Wickham et al., 2026), tidyr (Wickham et al., 2024), and networkD (Allaire et al., 2025). Habitat suitability dynamics were depicted using faceted stacked area plots partitioning gain, loss, and stable areas across time periods and SSP scenarios, while variable contributions were represented through Sankey diagrams to illustrate proportional influence on species distributions. To test our hypothesis (H4), where we predicted that areas characterized by moderate temperatures, higher dry-season precipitation, north-facing aspects, and gentle slopes would function as stable refugia for both species, whereas extreme warming, greater temperature variability, steep slopes, and south-facing aspects would reduce habitat suitability and promote fragmentation, we assessed the influence of these environmental and topographic factors on species distributions.

### 2.9 Model performance

Model evaluation indicated reliable predictive performance for both species, with clearer discrimination achieved for *Gentiana kurroo*. For *Gentiana cachemirica*, the model achieved a mean sensitivity of 0.72 and specificity of 0.83, resulting in a moderate mean TSS of 0.55. Discriminatory ability was supported by a high mean training AUC (0.93) and a good mean test AUC (0.80), signifying satisfactory generalization. The optimal probability threshold was 0.16, indicating a relatively permissive presence classification. In contrast, the *G. kurroo* model showed consistently stronger performance, with high mean sensitivity (0.82) and specificity (0.84) and a higher mean TSS of 0.67. Discrimination was strong, as reflected by a mean training AUC of 0.97 and a mean test AUC of 0.87. The lower optimal threshold (0.06) indicated a higher sensitivity to suitable conditions. Model performance therefore captured environmental responses of *G. kurroo* more clearly than those of *G. cachemirica*.

### 2.10 Hypothesis testing

To quantify the potential projected habitat contraction of *Gentiana cachemirica*, we used regression-based trend analyses (James et al., 2021). Specifically, we tested the prediction that (H1) increasing maximum temperature of the warmest month and reduced precipitation of the driest month shrink and fragment habitat, restricting the species to high-elevation, north-facing refugia. To do so, we fitted ordinary least squares regression models with habitat area (km²) as a function of time. We evaluated trend strength using the regression slope (*β*), coefficient of determination (R²), and p-values. To do so, we calculated percentage change relative to baseline conditions for each scenario. We also fitted scenario-specific regressions to assess directional consistency across climate pathways.

We evaluated scenario-driven habitat responses in *Gentiana kurroo* using non-parametric comparisons. Here, we tested whether (H2) *G. kurroo* persists or expands under low-emission scenarios by tracking suitable climates, but declines under stronger warming due to increasing temperature, variability, and drier conditions. To that end, we assessed normality of projected habitat-area values using the Shapiro–Wilk test and applied a Kruskal–Wallis test to compare habitat area across the different scenarios. To do so, we performed pairwise Wilcoxon rank-sum tests with Bonferroni correction where required. Finally, we used a paired Wilcoxon signed-rank test to compare habitat gain and loss.

We compared spatial responses between species using centroid displacement analysis. We tested whether (H3) *G. cachemirica* shows limited range shifts and greater habitat loss, while *G. kurroo* shows larger and directional shifts toward higher elevations and east-southeast directions. To test this hypothesis, we calculated centroid displacement distances and directional bearings for each species using paired Wilcoxon signed-rank tests to compare movement magnitude between species. We selected non-parametric tests due to non-normal distributions.

We identified environmental drivers of habitat stability using regression-based analyses. Specifically, we tested whether (H4) moderate climatic conditions, higher dry-season precipitation, north-facing slopes, and gentle terrain define refugia, while extreme warming, high variability, steep slopes, and south-facing aspects drive habitat loss and fragmentation. To do so, we fitted regression models linking stable habitat area to climatic and topographic predictors. We first included temperature, precipitation, slope, and aspect as explanatory variables. Next, we evaluated predictor influence using standardized regression coefficients (*β*). We quantified relative variable importance using permutation-based importance metrics, calculated as the reduction in model performance (AUC) following random permutation of predictor variables (Fisher et al., 2019).

## 3 Results

### 3.1 Effects of climate change on *Gentiana cachemirica*

Our models suggest that *Gentiana cachemirica* may face substantial habitat contraction under future climate change relative to baseline conditions (Fig. 1A; Fig. 2A–D; Fig. 3A–B). Supporting H1, increasing maximum temperature of the warmest month and reduced precipitation of the driest month are projected to shrink and fragment suitable habitat, restricting populations to high-elevation refugia (Fig. 2A–D). Suitable habitat is projected to decline from the current extent of 651 km² under all SSP scenarios. Under SSP126, habitat suitability is projected to decline by −37.5% during 2041–2060 and by −56.8% during 2081–2100. Stronger reductions are projected under SSP245 and SSP370, reaching −69.1% and −69.7%, respectively, by the end of the century. SSP585 is projected to produce the greatest decline, with habitat loss peaking at −72.2% during 2061–2080, followed by a partial recovery to −55.3% during 2081–2100 (Fig. 3A–B). Global regression analysis indicates a significant loss of suitable habitat of 4.35 km² per year (R² = 0.555, P = 0.0035). Scenario-specific regressions consistently produced negative slopes, although relationships remained non-significant (P > 0.05) because of the limited number of temporal projections (n = 3). Explanatory power was highest under SSP245 (R² = 0.872) and lowest under SSP585 (R² = 0.149) (Table S1–S3). Overall, projected habitat contraction is expected to increasingly confine *G. cachemirica* to fragmented high-elevation refugia across all scenarios (Fig. 2A–D; Fig. S1A–C to Fig. S4A–C).

**Figure 2.**
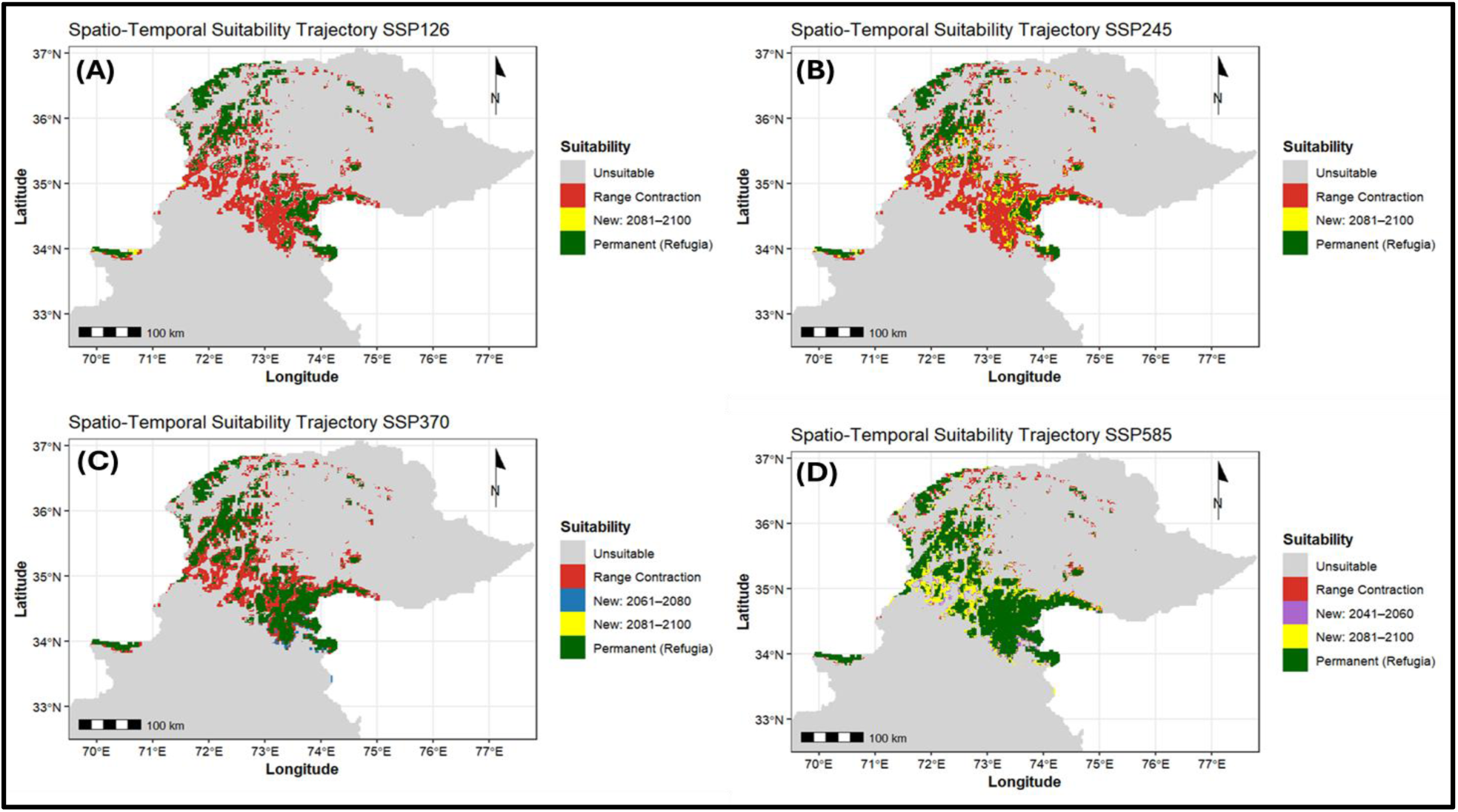
Projected spatio-temporal changes in habitat suitability of *Gentiana cachemirica* under future climate scenarios. Panels show projected changes in habitat suitability under the shared socioeconomic pathway scenarios (A) SSP126, (B) SSP245, (C) SSP370, and (D) SSP585. The maps illustrate the spatial distribution of unsuitable areas (grey), projected range contraction relative to baseline conditions (red), newly suitable habitats emerging during future periods (blue and yellow), and climatically stable refugia that remain suitable across all periods (green). Geographic coordinates are shown in latitude and longitude, and the scale bar represents 100 km. Map (A) shows projected habitat trajectories under SSP126, where habitat contraction occurs primarily in lower-elevation regions, while stable refugia remain concentrated in northern and high-elevation areas. Limited newly suitable habitats emerge during 2081–2100, although these areas remain spatially restricted. Map (B) shows stronger habitat contraction under SSP245, with increasing fragmentation across central and southern regions. Small patches of newly suitable habitat emerge during 2081–2100, although stable refugia remain confined to fragmented high-elevation zones. Map (C) shows continued habitat contraction under SSP370, with substantial fragmentation across the projected distribution range. Newly suitable habitats emerge during 2061–2080 and 2081–2100, although these areas remain scattered and disconnected. Stable refugia persist mainly within northern mountainous regions. Map (D) shows projected habitat changes under SSP585, where projected warming produces extensive habitat contraction across much of the current range. Although larger areas of newly suitable habitat emerge during future periods, these habitats remain fragmented and geographically restricted.

**Figure 3.**
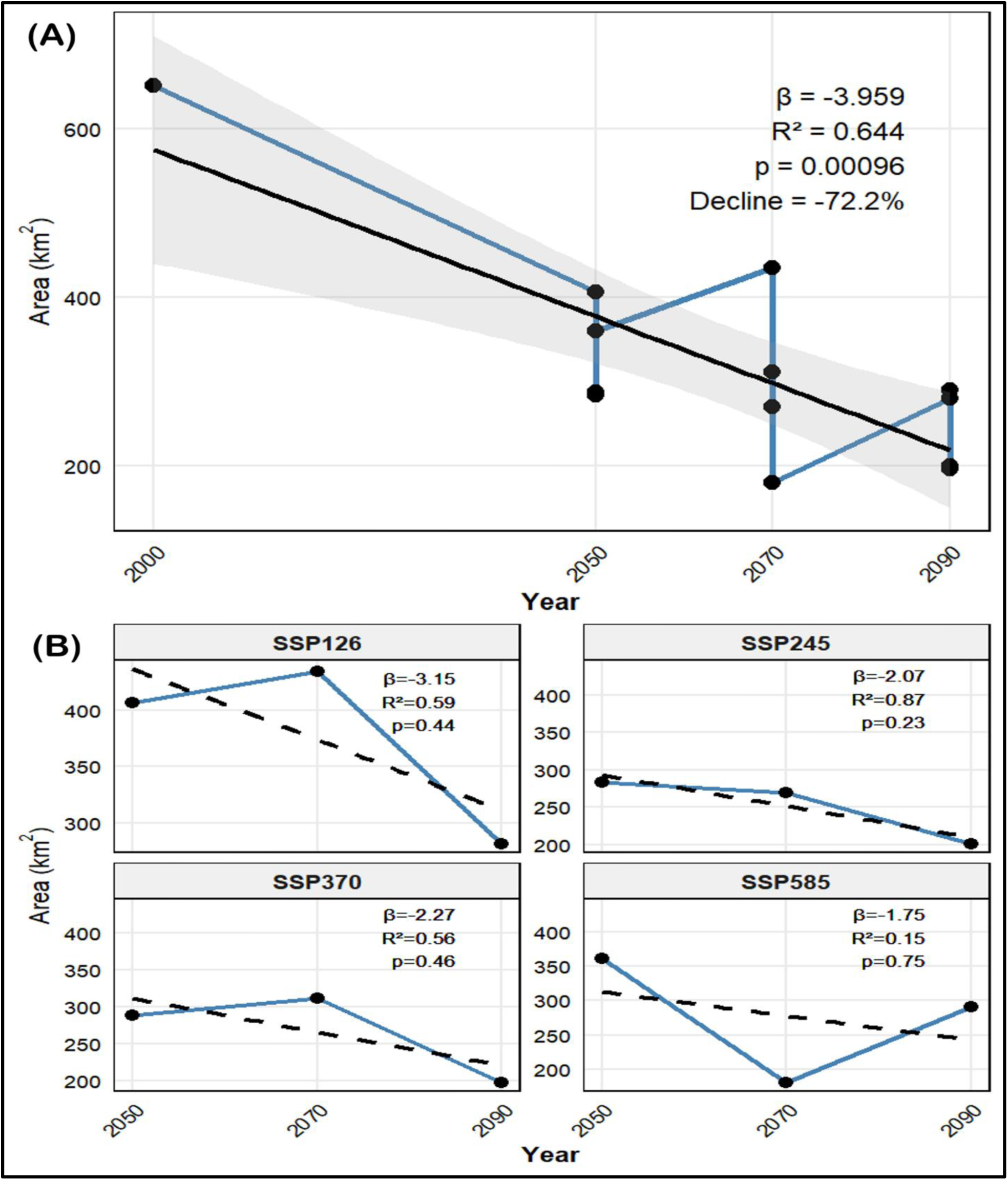
*Gentiana cachemirica* is projected to experience progressive range contraction across all future climate scenarios, with habitat loss being most pronounced under SSP585. (A) shows the overall temporal trend in suitable habitat area from the baseline period (2000) to future projection periods (2050, 2070, and 2090). The solid blue line represents projected changes in suitable habitat area across time, whereas the solid black regression line indicates the overall temporal trend. Shaded grey regions represent the confidence interval around the regression estimate. Global regression analysis indicates a significant negative relationship between time and habitat area (*β* = −3.96 km² yr⁻¹, R² = 0.644, P = 0.00096), with projected habitat loss reaching a maximum decline of −72.2% relative to baseline conditions. (B) shows scenario-specific temporal trends in suitable habitat area under the shared socioeconomic pathway scenarios SSP126, SSP245, SSP370, and SSP585. Blue solid lines represent projected habitat changes between future periods, whereas black dashed lines represent fitted linear regression trends within each scenario. All future climate scenarios show negative regression slopes, indicating continued habitat contraction through time. SSP126 projects an initial increase in suitable habitat between 2050 and 2070, followed by a decline by 2090. SSP245 and SSP370 show relatively consistent reductions in habitat extent across future periods. SSP585 projects the greatest variability, with a pronounced decline by 2070 followed by partial recovery by 2090.

### 3.2 Habitat dynamics of *Gentiana kurroo*

Projected habitat dynamics for *Gentiana kurroo* vary across climate scenarios and future periods (Fig. 1B, D; Fig. 4; Fig. 5A–D). We found partial support for H2, with *G. kurroo* projected to persist or locally expand under low-emission scenarios by tracking suitable climates toward higher elevations and east-southeast regions. However, stronger warming is projected to reduce habitat suitability and increase fragmentation through rising temperatures, greater climatic variability, and drier microclimatic conditions. By the end of the century, mean suitable habitat area is projected to decline to 1,436 km² ± 684.18 (S.D.), representing an average reduction of −41.44% ± 27.9 (S.D.) relative to baseline conditions. Net habitat change is projected to remain negative across all scenarios (*μ* = −1,010 km² ± 684.18, with habitat loss consistently exceeding habitat gain. Under low-emission scenarios, early future periods are projected to show localized habitat expansion, whereas stronger warming is expected to drive continued contraction during later periods (Fig. 5A–D; Table S1). The Kruskal–Wallis test detected no significant differences among climate scenarios (*χ²* = 11, df = 11, P = 0.443), although the Wilcoxon signed-rank test showed a significant difference between habitat gain and loss (V = 3, P = 0.002; Table 1). Overall, persistent negative habitat change indicates that projected warming may increasingly constrain long-term habitat availability for *G. kurroo* (Fig. 4). Supplementary projections for SSP126, SSP245, SSP370, and SSP585 are presented in Fig. S1D–F, Fig. S2D–F, Fig. S3D–F, and Fig. S4D–F, respectively.

**Figure 4.**
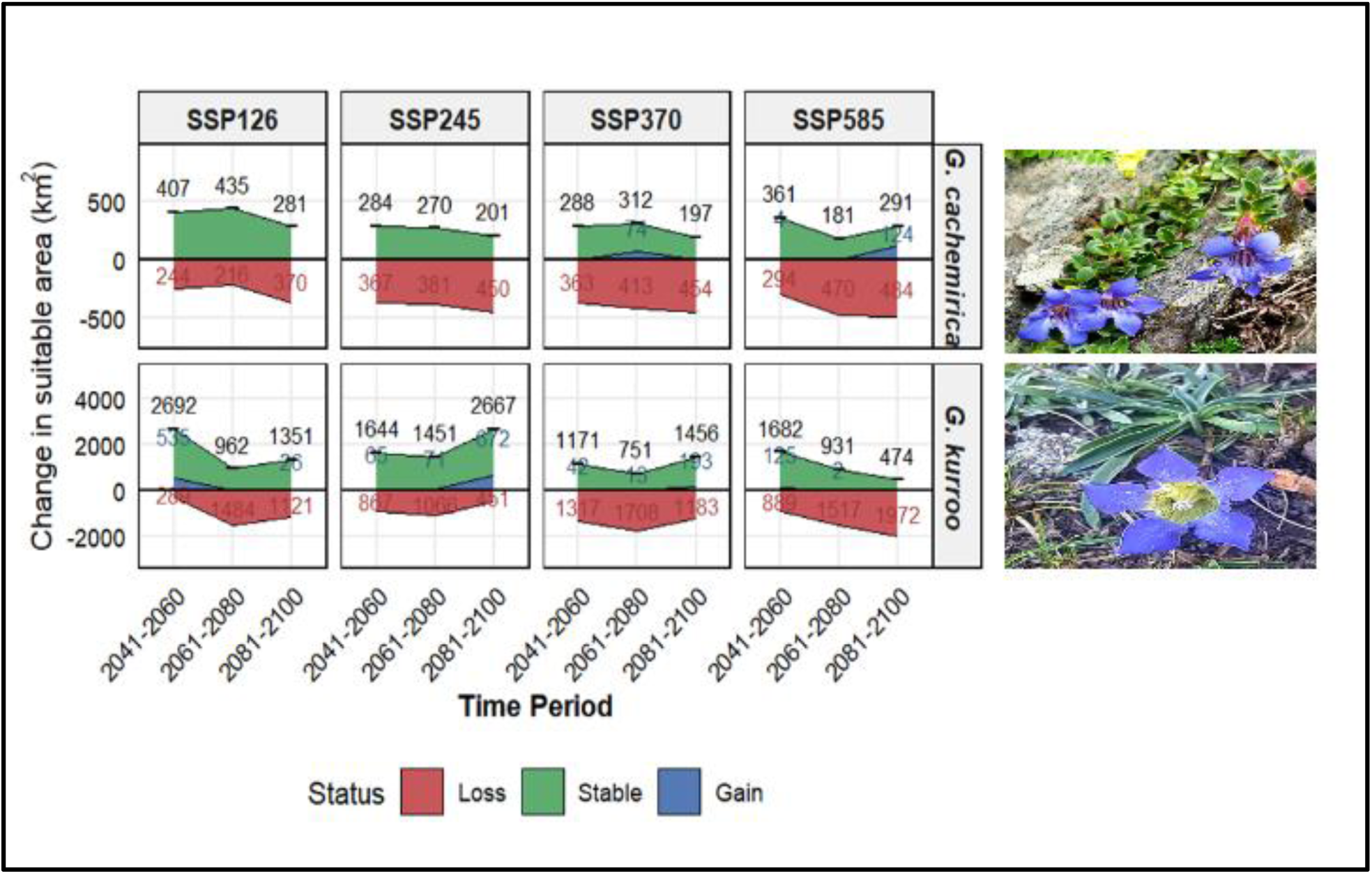
Future projections show increasing habitat loss and declining stability, with *Gentiana cachemirica* more strongly affected than *G. kurroo*. Predicted changes in suitable niche area (km²) are shown for both species across four Shared Socioeconomic Pathways (SSP126, SSP245, SSP370, and SSP585) and three future time periods (2041–2060, 2061–2080, and 2081–2100). Suitable areas are divided into habitat loss (red), stable habitat (green), and habitat gain (blue). Numerical values indicate the area (km²) for each category within each period. For *G. cachemirica* (top panels), stable habitat decreases with increasing emissions, alongside rising losses and limited gains. For *G. kurroo* (bottom panels), larger areas remain stable, although losses increase and gains vary across the different climatic scenarios.

**Figure 5.**
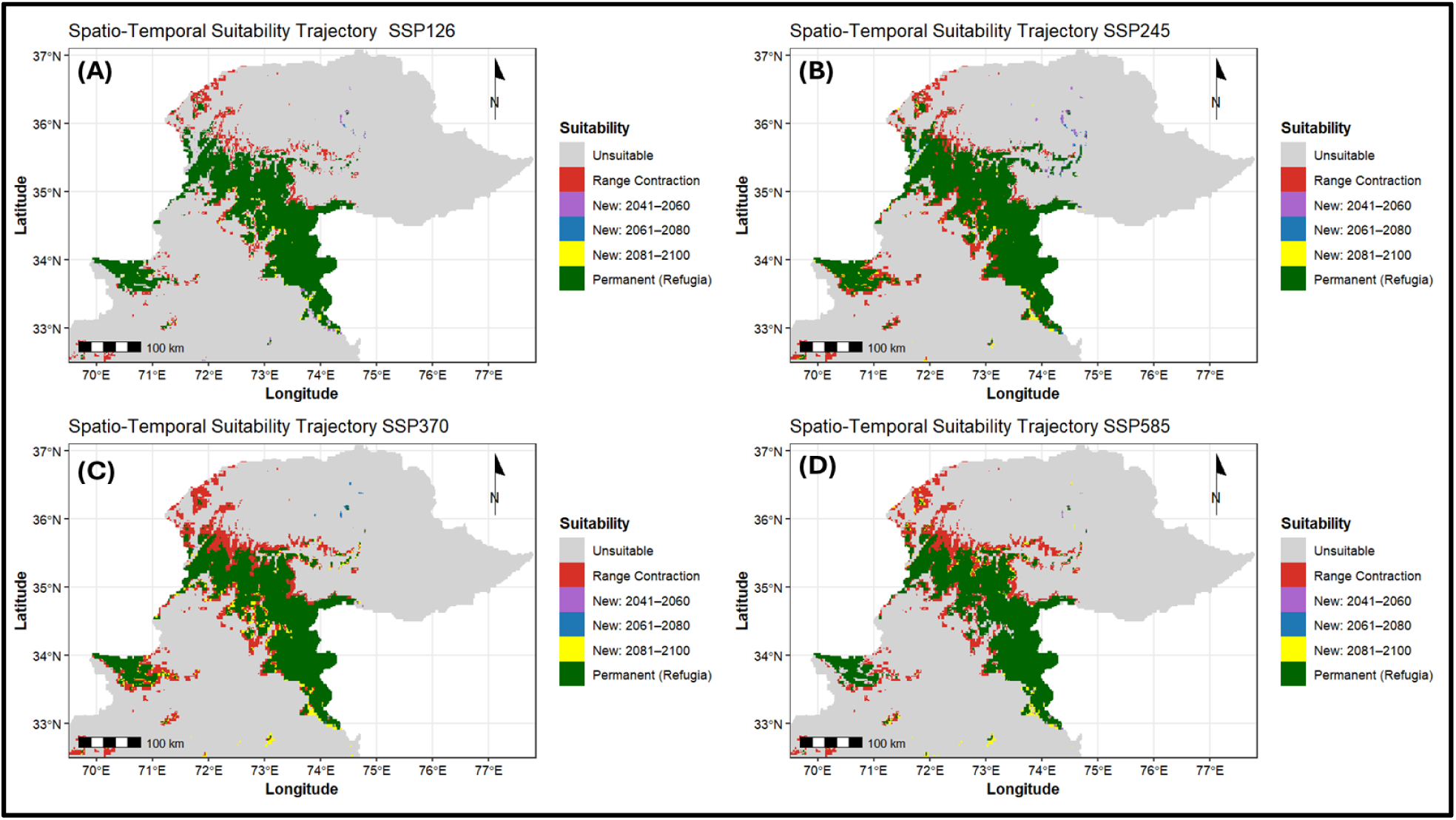
Projected habitat contraction and fragmentation increase with warming in *Gentiana kurroo*, despite the persistence of climatically stable refugia across all scenarios. Maps (A–D) show projected changes in habitat suitability under the shared socioeconomic pathway scenarios SSP126, SSP245, SSP370, and SSP585, respectively. Maps illustrate the spatial distribution of unsuitable areas (grey), projected range contraction relative to baseline conditions (red), newly suitable habitats emerging during 2041–2060 (purple), 2061–2080 (blue), and 2081–2100 (yellow), and climatically stable refugia that remain suitable across all periods (green). Geographic coordinates are shown in latitude and longitude, and the scale bar represents 100 km. Map (A) shows projected habitat trajectories under SSP126, where broad and relatively continuous refugial zones persist across central and southern regions. Habitat contraction remains concentrated around peripheral areas, whereas newly suitable habitats emerge in small and spatially restricted patches. Map (B) shows stronger habitat contraction under SSP245, with increasing fragmentation across northern and western regions. Stable refugia remain extensive, although newly suitable habitats become more scattered across future periods. Map (C) shows continued habitat contraction under SSP370, with increasing fragmentation and reduction of suitable habitat continuity. Newly suitable habitats emerge during later future periods, although these areas remain limited and disconnected. Map (D) shows habitat trajectories under SSP585, where stronger climate forcing produces substantial habitat contraction across the projected range. Stable refugial zones persist mainly within central mountainous regions, whereas newly suitable habitats remain fragmented and geographically restricted. Across all climate scenarios, *G. kurroo* maintains broader and more continuous refugial zones than *G. cachemirica*, although increasing warming intensifies habitat contraction and spatial fragmentation, particularly under SSP370 and SSP585.

**Table 1.** Projected habitat loss exceeds habitat gain in *Gentiana kurroo* across future climate scenarios. The table summarizes descriptive statistics and non-parametric test results associated with changes in suitable habitat area, including mean habitat area, percentage habitat change, and net habitat change relative to baseline conditions. Statistical analyses include the Shapiro–Wilk normality test, Kruskal–Wallis group comparison among climate scenarios, and Wilcoxon signed-rank test comparing habitat gain and habitat loss. Abbreviations are as follows: SD – standard deviation; W – Shapiro–Wilk test statistic; *χ²* – Kruskal–Wallis test statistic; V – Wilcoxon signed-rank statistic.

| No. | Category | Metric / test | Statistic | P-value |
| --- | --- | --- | --- | --- |
| 1 | Summary statistics | Mean area (km<sup>2</sup>) (± S.D.) | 1,436.00 ± 684.18 | |
| | | Mean % change (± S.D.) | –41.44 ± 27.90 | |
| | | Mean net change (km<sup>2</sup>) (± S.D.) | –1010.00 ± 684.18 | |
| 2 | Normality test | Shapiro–Wilk | W = 0.911 | 0.217 |
| 3 | Group comparison | Kruskal–Wallis | $\chi^2 = 11$ | 0.443 |
| 4 | Gain vs. loss | Wilcoxon signed-rank test | V = 3 | 0.002 |

### 3.3 Contrasting responses between species

Our projections suggest that *Gentiana kurroo* and *G. cachemirica* may undergo contrasting range shifts under future climate conditions (Fig. 2A–D; Fig. 5A–D). We found strong support for H3, where we predicted that *G. cachemirica* would show limited range shifts and greater habitat loss because of its narrow niche. In comparison, in our models, *G. kurroo* is projected to exhibit larger directional shifts toward higher elevations and east-southeast regions. Broader niche breadth and greater dispersal capacity likely support these spatial responses. *Gentiana kurroo* is projected to show a significantly greater mean shift distance (23.49 km) than *G. cachemirica* (12.72 km; Wilcoxon test: V = 12, P = 0.034). Our models further predict directional shifts to differ significantly between species. Indeed, *G. kurroo* is expected to show a mean bearing of 131.41°, indicating movement toward east-southeast regions, whereas *G. cachemirica* is expected to display a 24.88° bearing (V = 3, P = 0.005). These projected range shifts suggest greater spatial adjustment in *G. kurroo* under future climate conditions. By contrast, *G. cachemirica* may experience limited movement capacity and stronger habitat restriction across future scenarios (Fig. 4). Further details regarding centroid positions, migration distances, and shift directions are presented in Supplementary Material (Table S4 & S5).

### 3.4 Environmental drivers of habitat stability and loss

Our analyses indicate that climatic and topographic variables may strongly influence habitat stability and habitat loss in both species (Table 2). Specifically, we found strong support for H4, where we predicted that moderate temperatures, higher dry-season precipitation, north-facing slopes, and gentle terrain would support stable refugia. In contrast, our models predict that warming, high climatic variability, steep slopes, and south-facing aspects may increase habitat loss and fragmentation. For *G. cachemirica*, precipitation of the driest month (BIO14; *β* = 1.28, importance = 0.225) is expected to exert the strongest positive effect on habitat suitability, indicating that higher dry-season moisture may maintain habitat stability. Similarly, maximum temperature of the warmest month (BIO5; *β* = 0.831, importance = 0.115) and slope (*β* = 0.498, importance = 0.040) are likely to reduce habitat suitability, whereas aspect may exert a weak positive influence associated with north-facing slopes. For *G. kurroo*, temperature annual range (BIO7; *β* = 2.408, importance = 0.445) is predicted to exert the strongest influence on habitat suitability, indicating greater sensitivity to climatic variability. Maximum temperature and slope are also expected to reduce habitat suitability. Aspect may produce weak and inconsistent effects, whereas precipitation seasonality (BIO15) is likely to contribute only slightly to suitability patterns (Table 2).

**Table 2.** Temperature and precipitation dominate projected habitat suitability, whereas slope and aspect contribute to habitat stability and refugia in *Gentiana cachemirica* and *G. kurroo.* Effect strength coefficients (*β*), permutation importance values, direction of effect, and ecological interpretation for climatic and topographic variables included in habitat suitability models. Positive effect values indicate environmental conditions associated with increased habitat suitability or climatic refugia, whereas negative effect values indicate conditions associated with habitat decline or fragmentation. Abbreviations are as follows: BIO5 – maximum temperature of the warmest month; BIO7 – annual temperature range; BIO14 – precipitation of the driest month; BIO15 – precipitation seasonality; *β* – effect strength coefficient.

| Species | Variable | Effect strength ($\beta$) | Permutation importance | Effect direction | Interpretation |
| --- | --- | --- | --- | --- | --- |
| *Gentiana cachemirica* | BIO14 | 1.280 | 0.225 | Positive | Higher dry-season precipitation supports refugia |
|  | BIO5 | 0.831 | 0.115 | Negative | Increasing temperature reduces suitability |
|  | Slope | 0.498 | 0.04 | Negative | Steep terrain reduces stability |
|  | Aspect | 0.094 | 0.004 | Positive (north) | North-facing slopes favour refugia |
| *Gentiana kurroo* | BIO7 | 2.408 | 0.445 | Negative | High temperature variability drives habitat loss |
|  | BIO5 | 0.258 | 0.005 | Negative | Warming negatively affects the habitat |
|  | Slope | 0.355 | 0.008 | Negative | Steep slopes increase fragmentation |
|  | Aspect | 0.357 | -0.005 | Weak/inconsistent | Aspect has limited influence |
|  | BIO15 | 0.094 | 0.001 | Weak positive | Minor role of precipitation variability |

## 4. Discussion

Climate change has already caused a dent on mountain biodiversity (Salick et al., 2014; Dainese et al., 2024), particularly among narrowly endemic species (Manes et al., 2021). This loss is critical, as mountain biodiversity supports ecosystem stability, maintains endemic species, and regulates water resources under changing climatic conditions (Spehn et al., 2011; Perrigo et al., 2020). Here, we examined the future climatic impacts on the habitat suitability of two endemic and endangered *Gentiana* species in the Western Himalayas. We built species distribution models to project habitat suitability, range shifts, and examine the key climatic determinants under multiple emission scenarios. We tested whether climate warming restricts and fragments the habitat of *G. cachemirica* and whether *G. kurroo* maintains temporary persistence under low-emission scenarios but declines under stronger warming. We also tested whether both species differ in range shifts, climatic sensitivity, and refugial stability because of variation in niche breadth and dispersal capacity. Our approach provides a spatially explicit assessment of climate-driven redistribution across elevational gradients. Our models predict that *G. cachemirica* may undergo consistent habitat contraction across all climate scenarios, indicating high vulnerability to warming. By contrast, *G. kurroo* may exhibit limited habitat expansion under low-emission conditions but may decline under moderate- and high-warming scenarios. Both species are projected to exhibit contrasting ecological sensitivities and spatial responses to climatic predictors, consistent with reported alpine plant range shifts (Chen et al., 2011; Lamprecht et al., 2018). Our findings align with Dagnino et al. (2017), highlighting the importance of differences in ecological niche structure and climatic tolerance in driving divergence in habitat suitability and spatial distribution patterns under changing environmental conditions.

Endemic species are more vulnerable to climate change than widespread species because endemic species’ narrow ecological ranges typically limit their adaptive potential (Manzoor et al., 2026). Here, our projections suggest that *G. cachemirica*, a range-restricted endemic Himalayan species, may undergo strong habitat contraction and increasing fragmentation under all future climate scenarios we examined. According to our forecasts, its suitable habitat is expected to become progressively restricted and ultimately isolated to high elevation refugia in the Himalayas. Our projections are consistent with previous studies suggesting that endemic mountain species with narrow climatic niches may be highly sensitive to warming-driven habitat loss and range fragmentation (Wershow & DeChaine, 2018; Khan et al., 2025). Such responses are consistent with the restricted climatic tolerance and sensitivity to temperature and moisture variability that characterize many alpine-endemic (Larcher et al., 2010; Körner, 2014). Habitat contraction of *G. cachemirica* is projected to intensify under our examined future scenarios, with no compensatory range expansion reported. As such, our projections further align with studies reporting upward range shifts, habitat compression, and declining connectivity among alpine plant populations under climate warming (Dirnböck et al., 2011; Dullinger et al., 2012; Pauli & Halloy, 2019). The resulting reduction of suitable habitat to increasingly isolated patches may elevate extinction risk by reducing dispersal opportunities and increasing population isolation (Niebuhr et al., 2015).

Climate change is affecting the distribution and persistence of threatened mountainous flora, highlighting the need for effective conservation and adaptive management strategies in mountain ecosystems (Körner, 2014; Gillani et al., 2026). In our case, our projections suggest that *Gentiana kurroo*, a critically endangered species, may show partial resilience under low-emission climate scenarios, followed by pronounced decline under stronger warming conditions. A key prediction of our models is that *G. kurroo* occupies a broader and more continuous baseline distribution than *G. cachemirica*, consistent with its wider climatic tolerance and greater ecological flexibility across mid-elevation habitats. Indeed, under the SSP126 and SSP245 climatic scenarios, the suitable habitat of this species is projected to increase during mid-century, reflecting short-term climatic alignment and upslope redistribution into newly suitable areas. Here, climatic suitability is projected to show significant temporal instability, with initial habitat gains not sustained in later periods, indicating a non-persistent response under future climate scenarios. Under the SSP370 and SSP585 scenarios, our projections suggest that the suitable habitat of *G. kurroo* may decline sharply, reaching approximately one-fifth of baseline extent by the end of the century. Our projections are consistent with those reported by Pauli & Halloy (2019), who found that climate warming can drive upward range shifts while reducing the availability of suitable habitat for cold-adapted mountain species. Such responses suggest that species with intermediate climatic tolerance may initially benefit from moderate environmental change but may experience a sharp decline once physiological limits are exceeded (Thuiller et al., 2005; Slatyer et al., 2013; Evans & Jacquemyn, 2022). Similar temporal patterns, characterized by early expansion followed by contraction, have also been reported in montane systems under progressive warming (Lloret et al., 2012; Olivares et al., 2015). *Gentiana kurroo* therefore may exhibit non-linear climate responses, where short-term gains may mask long-term vulnerability under high-emission pathways.

We found strong support for our prediction that contrasting climatic responses may emerge between *G. cachemirica* and *G. kurroo* under projected climate change scenarios. Specifically, our models predict that *G. cachemirica* may face range contraction, fragmentation, and limited spatial adjustment, reflecting restricted dispersal and strong local dependence. Our projections are consistent with the findings by Yu et al. (2017), who reported that narrow range alpine species experienced greater habitat loss and range contraction than widespread species because of higher climatic sensitivity. In contrast, *G. kurroo* is projected to exhibit greater spatial flexibility, consistent with Rather et al. (2021), who found that species with broader climatic niches were more capable of shifting their distributions in response to climate change. Centroid analyses further suggest limited spatial displacement in *G. cachemirica*, indicating restricted redistribution potential under future climate change. This pattern is consistent with the projected contraction and fragmentation of suitable habitat and likely reflects limited capacity to track shifting climatic conditions. Similar responses were reported by Chen et al. (2026), who found that species with narrower ecological niches exhibited greater climatic sensitivity and more constrained distributional responses, and by Peña et al. (2024), who showed that dispersal limitations can reduce connectivity and restrict species movements across mountain landscapes. By contrast, *G. kurroo* is projected to exhibit more pronounced directional shifts, suggesting a greater capacity to adjust its distribution in response to changing climatic conditions. This pattern aligns with Littlefield et al. (2019), who emphasized that successful climate-driven range shifts depend on maintaining connectivity between current and future climatically suitable habitats. Here, our projections suggest that contrasting responses reflect differences in niche breadth and dispersal capacity, with narrow-niche taxa being more constrained and broader-niche taxa more responsive under climate change (Qiao et al., 2016; Saupe et al., 2015; Lenoir & Svenning, 2015).

Climatic and topographic gradients play a central role in shaping the distribution, stability, and persistence of mountain plant species under climate change (Sang, 2009; Manzoor et al., 2025). Here, our projections indicate that climatic and topographic variables strongly regulate habitat stability and patterns of habitat loss in both *Gentiana* species under future climate conditions. Temperature and precipitation are projected to remain the primary determinants of habitat suitability. Similar patterns have been reported, with these climatic variables strongly influencing species niches and distributional responses to environmental change (Elith & Leathwick, 2009; Franklin, 2023). Our analysis suggests that *G. cachemirica* is likely to remain particularly sensitive to precipitation of the driest month and maximum temperature, indicating strong dependence on moisture availability and thermal limits. By contrast, *G. kurroo* is projected to respond primarily to annual temperature range, suggesting greater sensitivity to seasonal climatic variability. Together, these projections suggest that stable habitats are most likely to persist in climatically buffered environments, particularly at higher elevations and on north-facing slopes (Singh, 2018). Habitat loss, however, is projected to increase under warmer conditions, increased climatic variability, steep slopes, and south-facing aspects that intensify heat exposure and reduce soil moisture. Similar patterns were reported by Ebel (2012), who found that warmer and drier conditions reduce habitat suitability. Our projections therefore align with previous studies showing that environmental gradients structure the spatial distribution of refugia and zones of decline across mountainous landscapes (Dobrowski, 2011; Scherrer & Körner, 2011; Gentili et al., 2015b). Such patterns are consistent with Graae et al. (2018), who highlighted the importance of topographic heterogeneity and microclimatic refugia in supporting species persistence under climate change.

Mountain ecosystems support high levels of biodiversity and endemism but remain highly vulnerable to ongoing climate change and habitat fragmentation (Gillani et al., 2026; Manzoor et al., 2026). The Himalayas are a biodiversity hotspot shaped by climatic and topographic gradients, supporting high habitat diversity (Trew & Maclean, 2021). Endemic species dominate Himalayan flora and contribute substantially to ecosystem structure, ecological functioning, and regional biodiversity patterns (Manzoor et al., 2026). Endemic species are also highly sensitive to environmental change, making them important indicators of ecosystem vulnerability (Manzoor et al., 2026). As such, the effective conservation of endemic Himalayan flora requires understanding how climate and landscape regulate species persistence. In this context, the contrasting responses of *G. cachemirica* and *G. kurroo* projected here to highlight the need to integrate climatic projections with dispersal constraints in conservation planning. Process-based and hybrid models that combine distribution, demography, and dispersal can improve predictions of future population viability (Dormann et al., 2012; Singer et al., 2018). However, limited field data on population dynamics, reproductive traits, and dispersal capacity remain critical knowledge gaps for alpine endemic species (Körner, 2014; Manzoor et al., 2026). We argue that long-term ecological monitoring across elevational gradients will remain essential to validate projected range shifts and detect early population declines. We also suggest that the protection of high elevation refugia should be prioritized to conserve climatically stable habitats expected to support vulnerable species (Khan et al., 2025; Körner, 2009). The maintenance of landscape connectivity along elevational gradients is expected to facilitate climate-driven species redistribution (Lenoir & Svenning, 2015; Guo et al., 2018). Finally, we recommend that integrating remote sensing, field surveys, and climate projections will improve the detection of habitat change and inform conservation actions (Rose et al., 2015).

## 5. Conclusions

Climate change increasingly threatens the persistence, distribution, and ecological stability of mountain biodiversity globally. Climate change poses a significant threat to Himalayan *Gentiana* species, with clear interspecific differences in vulnerability. Here, our projections indicate that the endemic Himalayan species *Gentiana cachemirica* may experience consistent habitat contraction, increasing fragmentation, and limited range shifts across all climate scenarios, reflecting its narrow ecological niche and high climatic sensitivity. However, the broader-niche species *G. kurroo* is projected to exhibit partial resilience under low-emission conditions but is expected to undergo substantial declines under stronger warming, indicating that broader ecological tolerance does not fully buffer against climate stress. Projected habitat loss and directional range shifts highlight critical risks for mountain biodiversity, particularly for species with restricted distributions. High elevation refugia and climatically stable areas identified in our study represent key conservation priorities, as they are expected to provide significant buffers against ongoing environmental change. We argue that maintaining landscape connectivity will be crucial to facilitate species movement and reduce long-term extinction risk. Ultimately, integrating climate projections into conservation planning, along with habitat protection and ecological monitoring, will be essential for sustaining these species under future climatic conditions.

## Author contributions

SWG, MM, and RSG conceived the foundational ideas. SWG designed the methodology and conceptualized the study with input from RSG, MM and RWAK. SWG, MM and AS sourced and collected the filed data. RWAK lead the analysis and data visualization with input from SWG and MM. MA and RSG supervised the study. RSG and SWG developed the hypotheses. RSG analytically reviewed and improved the initial draft. SWG led the first draft of the manuscript under the guidance of RSG. All authors edited the draft for final submission.

## Acknowledgments

We acknowledge the Department of Plant Sciences, Quaid-i-Azam University, Islamabad, Pakistan, and the Department of Biology, University of Oxford, United Kingdom, for providing support during initial data collection and access to facilities for manuscript completion. SWG was supported by a Commonwealth Split-Site Scholarship at Oxford. RSG was supported by a NERC Pushing the Frontiers grant (NE/X013766/1). We thank the local communities of the Himalayan region for their invaluable assistance during fieldwork.

