## Supplementary tables for "Climate change drives contrasting redistribution patterns in endemic and endangered Himalayan *Gentiana*"

**Table S1. Scenario-wise projected changes in suitable habitat area of *Gentiana cachemirica* and *G. kurroo* under future climate scenarios.** The table presents total suitable habitat area (km²), habitat gain (km²), habitat loss (km²), and stable habitat area (km²) for both species across the shared socioeconomic pathway scenarios SSP126, SSP245, SSP370, and SSP585 relative to baseline climatic conditions. Habitat gain represents newly suitable areas projected under future climate conditions, habitat loss represents areas projected to become unsuitable relative to baseline conditions, and stable habitat represents climatically suitable areas maintained across present and future periods. Abbreviations are as follows: SSP – shared socioeconomic pathway; NA – not applicable.

| **Scenario** | **Suitable area (km²)** | | **Gain (km²)** | | **Loss (km²)** | | **Stable (km²)** | |
|---|---|---|---|---|---|---|---|---|
| | ***G. cachemirica*** | ***G. kurroo*** | ***G. cachemirica*** | ***G. kurroo*** | ***G. cachemirica*** | ***G. kurroo*** | ***G. cachemirica*** | ***G. kurroo*** |
| **Baseline** | 651 | 2452 | NA | NA | NA | NA | NA | NA |
| SSP126 (2041–2060) | 407 | 2692 | 0 | 535 | 244 | 289 | 407 | 2157 |
| SSP126 (2061–2080) | 435 | 962 | 0 | 0 | 216 | 1484 | 435 | 962 |
| SSP126 (2081–2100) | 281 | 1351 | 0 | 26 | 370 | 1121 | 281 | 1325 |
| SSP245 (2041–2060) | 284 | 1644 | 0 | 65 | 367 | 867 | 284 | 1579 |
| SSP245 (2061–2080) | 270 | 1451 | 0 | 71 | 381 | 1066 | 270 | 1380 |

| | | | | | | | | |
|---|---|---|---|---|---|---|---|---|
| SSP245 (2081–2100) | 201 | 2667 | 0 | 672 | 450 | 451 | 201 | 1995 |
| SSP370 (2041–2060) | 288 | 1171 | 0 | 42 | 363 | 1317 | 288 | 1129 |
| SSP370 (2061–2080) | 312 | 751 | 74 | 13 | 413 | 1708 | 238 | 738 |
| SSP370 (2081–2100) | 197 | 1456 | 0 | 193 | 454 | 1183 | 197 | 1263 |
| SSP585 (2041–2060) | 361 | 1682 | 4 | 125 | 294 | 889 | 357 | 1557 |
| SSP585 (2061–2080) | 181 | 931 | 0 | 2 | 470 | 1517 | 181 | 929 |
| SSP585 (2081–2100) | 291 | 474 | 124 | 0 | 484 | 1972 | 167 | 474 |

**Table S2. Percentage change in suitable habitat area, net habitat change, and habitat stability of *Gentiana cachemirica* and *Gentiana kurroo* under future climate scenarios.** The table presents percentage change in suitable habitat area (Δ Area), net habitat change ($km^2$), and habitat stability (%) for both species across the shared socioeconomic pathway scenarios SSP126, SSP245, SSP370, and SSP585 during future projection periods relative to baseline habitat extent (651 $km^2$ for *G. cachemirica* and 2452 $km^2$ for *G. kurroo*). Negative percentage values and net change values indicate projected habitat contraction relative to baseline conditions, whereas habitat stability values represent the proportion of climatically suitable habitat maintained across present and future periods. Abbreviations are as follows: SSP – shared socioeconomic pathway; Δ Area – percentage change in suitable habitat area.

| Scenario | Δ Area (%) | | Net change ($km^2$) | | Stability (%) | |
|---|---|---|---|---|---|---|
| | *G. cachemirica* | *G. kurroo* | *G. cachemirica* | *G. kurroo* | *G. cachemirica* | *G. kurroo* |
| SSP126 (2041–2060) | -37.5 | 9.8 | -244 | 246 | 62.5 | 87.9 |
| SSP126 (2061–2080) | -33.2 | -60.8 | -216 | -1484 | 66.8 | 39.2 |
| SSP126 (2081–2100) | -56.8 | -44.9 | -370 | -1095 | 43.2 | 54 |
| SSP245 (2041–2060) | -56.4 | -33 | -367 | -802 | 43.6 | 64.4 |
| SSP245 (2061–2080) | -58.5 | -40.8 | -381 | -995 | 41.5 | 56.3 |
| SSP245 (2081–2100) | -69.1 | 8.8 | -450 | 221 | 30.9 | 81.2 |
| SSP370 (2041–2060) | -55.8 | -52.2 | -363 | -1275 | 44.2 | 46 |

| | | | | | | |
|---|---|---|---|---|---|---|
| SSP370 (2061–2080) | -52.1 | -69.4 | -339 | -1695 | 36.6 | 30.1 |
| SSP370 (2081–2100) | -69.7 | -40.6 | -454 | -990 | 30.3 | 51.6 |
| SSP585 (2041–2060) | -44.5 | -31.4 | -290 | -764 | 54.8 | 63.5 |
| SSP585 (2061–2080) | -72.2 | -62 | -470 | -1515 | 27.8 | 37.9 |
| SSP585 (2081–2100) | -55.3 | -80.7 | -360 | -1972 | 25.7 | 19.3 |

**Table S3. Projected changes in suitable habitat area of *Gentiana cachemirica* under future climate scenarios.** The table presents projected habitat area ($km^2$), percentage habitat change relative to baseline conditions, and temporal regression statistics across future periods under the shared socioeconomic pathway scenarios SSP126, SSP245, SSP370, and SSP585. Regression outputs include slope coefficients ($\beta$), coefficients of determination ($R^2$), and associated p-values describing temporal trends in projected habitat extent. Negative percentage values indicate projected habitat contraction relative to baseline conditions. Abbreviations are as follows: SSP – shared socioeconomic pathway; $\beta$ – regression slope coefficient; $R^2$ – coefficient of determination.

| Scenario | Area ($km^2$) | Year | Percent change (%) | $\beta$ ($km^2\ yr^{-1}$) | $R^2$ | p-value |
|---|---|---|---|---|---|---|
| Baseline | 651 | 2020 | 0.0 | – | – | – |
| SSP126 (2041–2060) | 407 | 2050 | −37.5 | −3.15 | 0.590 | 0.443 |
| SSP126 (2061–2080) | 435 | 2070 | −33.2 | −3.15 | 0.590 | 0.443 |
| SSP126 (2081–2100) | 281 | 2090 | −56.8 | −3.15 | 0.590 | 0.443 |
| SSP245 (2041–2060) | 284 | 2050 | −56.4 | −2.08 | 0.872 | 0.233 |
| SSP245 (2061–2080) | 270 | 2070 | −58.5 | −2.08 | 0.872 | 0.233 |
| SSP245 (2081–2100) | 201 | 2090 | −69.1 | −2.08 | 0.872 | 0.233 |

| | | | | | | |
|---|---|---|---|---|---|---|
| SSP370 (2041–2060) | 288 | 2050 | −55.8 | −2.28 | 0.563 | 0.460 |
| SSP370 (2061–2080) | 312 | 2070 | −52.1 | −2.28 | 0.563 | 0.460 |
| SSP370 (2081–2100) | 197 | 2090 | −69.7 | −2.28 | 0.563 | 0.460 |
| SSP585 (2041–2060) | 361 | 2050 | −44.5 | −1.75 | 0.149 | 0.748 |
| SSP585 (2061–2080) | 181 | 2070 | −72.2 | −1.75 | 0.149 | 0.748 |
| SSP585 (2081–2100) | 291 | 2090 | −55.3 | −1.75 | 0.149 | 0.748 |
| **Overall trend** | | | **Max decline: −72.2%** | −4.35 | 0.555 | 0.0035 |

**Table S4. Centroid coordinates, migration distance, and direction of projected range shifts for *Gentiana cachemirica* and *G. kurroo* under baseline and future SSP scenarios.** This table summarizes spatial shifts in species distributions under different climate scenarios. Baseline centroid positions represent current distribution centres. Future projections are based on SSP126, SSP245, SSP370, and SSP585 across three time periods (2041-2060, 2061-2080, and 2081-2100). Migration distance is expressed in kilometres (km). Direction is reported as bearing (°), indicating the orientation of range shifts.

| Scenario | Centroid Lon (°E) | | Centroid Lat (°N) | | Distance (km) | | Bearing (°) | |
|---|---|---|---|---|---|---|---|---|
| | ***G. cachemirica*** | ***G. kurroo*** | ***G. cachemirica*** | ***G. kurroo*** | ***G. cachemirica*** | ***G. kurroo*** | ***G. cachemirica*** | ***G. kurroo*** |
| Baseline | 71.9477 | 73.5749 | 36.3225 | 34.3126 | – | 0 | – | NA |
| SSP126 (2041–2060) | 71.9477 | 73.6876 | 36.3225 | 34.232 | 0 | 13.71 | NA | 130.69 |
| SSP126 (2061–2080) | 71.9771 | 73.7429 | 36.4114 | 34.2029 | 10.24 | 19.7 | 15 | 128.13 |
| SSP126 (2081–2100) | 71.9431 | 73.7389 | 36.4105 | 34.1834 | 9.8 | 20.85 | −2.41 | 133.44 |
| SSP245 (2041–2060) | 71.9799 | 73.7256 | 36.4406 | 34.2388 | 13.46 | 16.12 | 12.45 | 120.48 |
| SSP245 (2061–2080) | 71.9718 | 73.7811 | 36.431 | 34.1763 | 12.27 | 24.3 | 10.18 | 128.46 |
| SSP245 (2081–2100) | 71.9785 | 73.7311 | 36.4397 | 34.1803 | 13.33 | 20.58 | 12 | 135.5 |
| SSP370 (2041–2060) | 71.9152 | 73.7943 | 36.3931 | 34.1432 | 8.37 | 27.63 | −20.39 | 132.85 |
| SSP370 (2061–2080) | 71.9548 | 73.7987 | 36.417 | 34.1674 | 10.54 | 26.18 | 3.49 | 127.93 |

| | | | | | | | | |
|---|---|---|---|---|---|---|---|---|
| SSP370 (2081–2100) | 72.0398 | 73.7915 | 35.8361 | 34.1271 | 54.78 | 28.71 | 171.23 | 135.82 |
| SSP585 (2041–2060) | 71.9091 | 73.758 | 36.3891 | 34.1834 | 8.18 | 22.16 | −25.08 | 130.29 |
| SSP585 (2061–2080) | 71.9762 | 73.798 | 36.298 | 34.1273 | 3.74 | 29.11 | 136.74 | 134.93 |
| SSP585 (2081–2100) | 71.8917 | 73.8139 | 36.3773 | 34.0937 | 7.9 | 32.84 | −39.55 | 137.72 |

**Table S5. Paired Wilcoxon tests comparing projected range-shift distance and direction between *Gentiana cachemirica* and *Gentiana kurroo* under future climate scenarios.** The table presents mean displacement distance (km) and mean shift direction (°) for both species, together with results of paired Wilcoxon tests assessing interspecific differences in projected spatial responses to climate change. Abbreviations are as follows: V – Wilcoxon signed-rank statistic; ° – directional bearing in degrees.

| Variable | *Gentiana cachemirica* (Mean) | *Gentiana kurroo* (Mean) | Test (Wilcoxon) | Statistic (V) | p-value |
|---|---|---|---|---|---|
| Distance (km) | 12.72 | 23.49 | Paired Wilcoxon | 12 | 0.034 |
| Direction (°) | 24.88 | 131.41 | Paired Wilcoxon | 3 | 0.0049 |
