## Supplementary figures for "Climate change drives contrasting redistribution patterns in endemic and endangered Himalayan *Gentiana*"

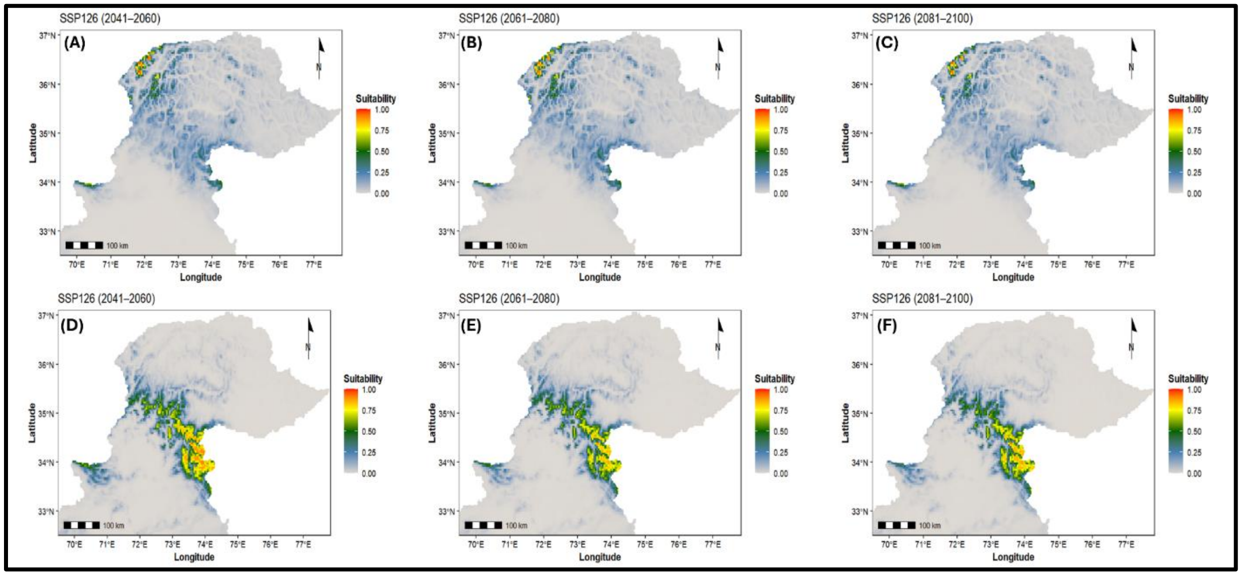


**Figure S1. Habitat suitability under SSP126 shows contraction in *Gentiana cachemirica* and a variable response in *G. kurroo.*** (A-C) Projected habitat suitability for *Gentiana cachemirica* across three future periods (2041-2060, 2061-2080, and 2081-2100), showing progressive reduction and increasing fragmentation of suitable areas, largely confined to higher elevations. (D-F) Corresponding projections for *G. kurroo*, illustrating an expansion of suitable habitat during mid-century, followed by reduction and spatial redistribution in later periods. Habitat suitability values range from 0 to 1 and are represented using a continuous color scale from low (blue) to high (red), derived from generalized linear models using projected climatic variables from the ACCESS-CM2 model. These patterns indicate that even under low-emission conditions, suitable habitat for *G. cachemirica* becomes increasingly limited, whereas *G. kurroo* exhibits short-term expansion that is not sustained over time.

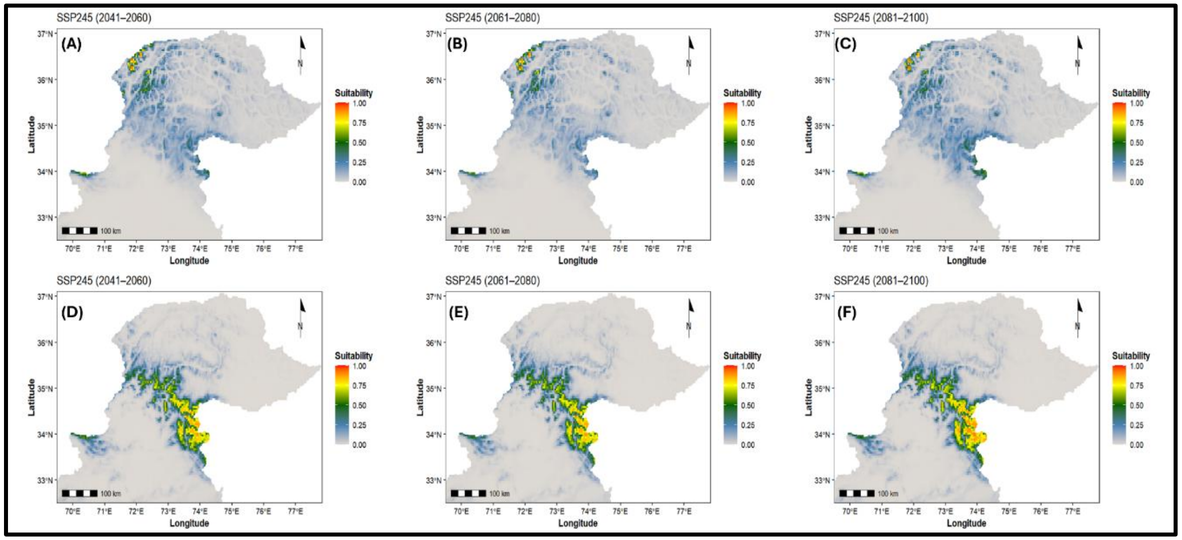


**Figure S2. Habitat suitability under SSP245 shows continued contraction in *Gentiana cachemirica* and a non-linear response in *G. kurroo*.** (A-C) Projected habitat suitability for *G. cachemirica* across three future periods (2041-2060, 2061-2080, and 2081-2100), showing steady reduction and increasing isolation of suitable habitat, while (D-F) corresponding projections for *G. kurroo* illustrate reduced suitability during mid-century followed by expansion toward the end of the century. Habitat suitability values range from 0 to 1 and are represented using a continuous color scale from low (blue) to high (red), derived from projected climatic conditions. These patterns indicate that moderate climate change leads to persistent habitat loss for *G. cachemirica*, whereas *G. kurroo* exhibits temporally variable suitability across future periods.

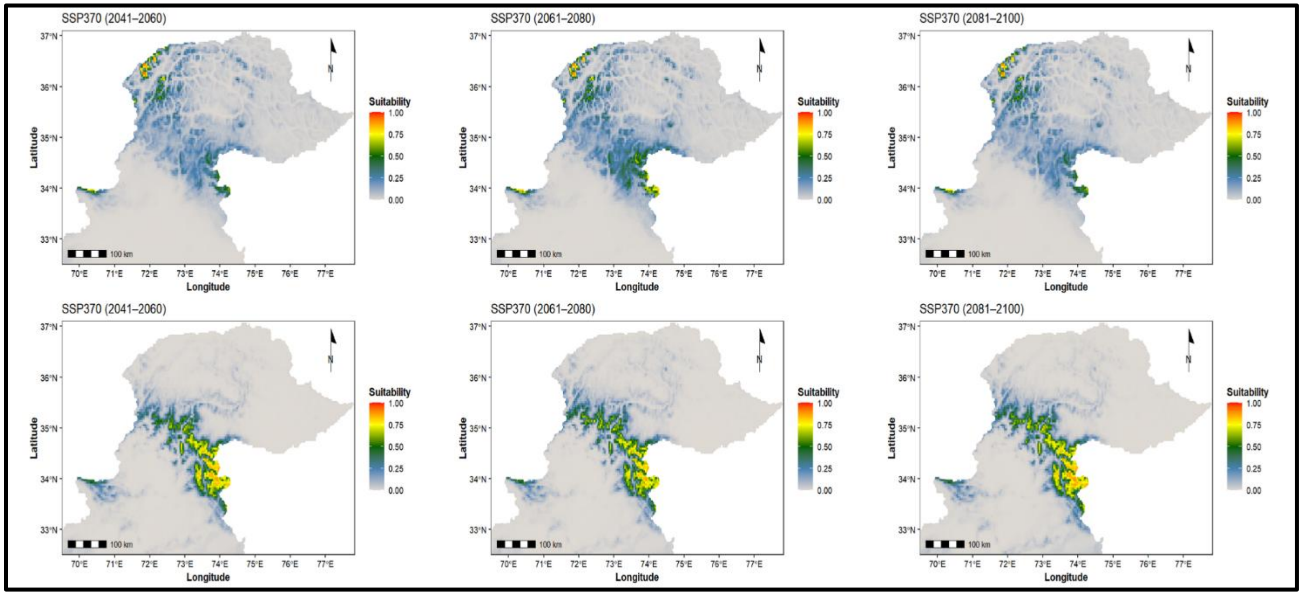


**Figure S3. Habitat suitability under SSP370 shows substantial reduction in suitable habitat for both species, indicating increasing climatic pressure**. (A-C) Projected suitability for *Gentiana cachemirica* shows strong contraction and restriction to high-altitude regions with limited continuity, while (D-F) projected suitability for *G. kurroo* indicates pronounced declines in suitable habitat with only minor recovery in later periods. Habitat suitability values (0-1) are derived from climate projections and represented using a continuous color gradient from low (blue) to high (red). These patterns indicate that higher levels of warming reduce both the extent and stability of suitable habitats for the two species.

.

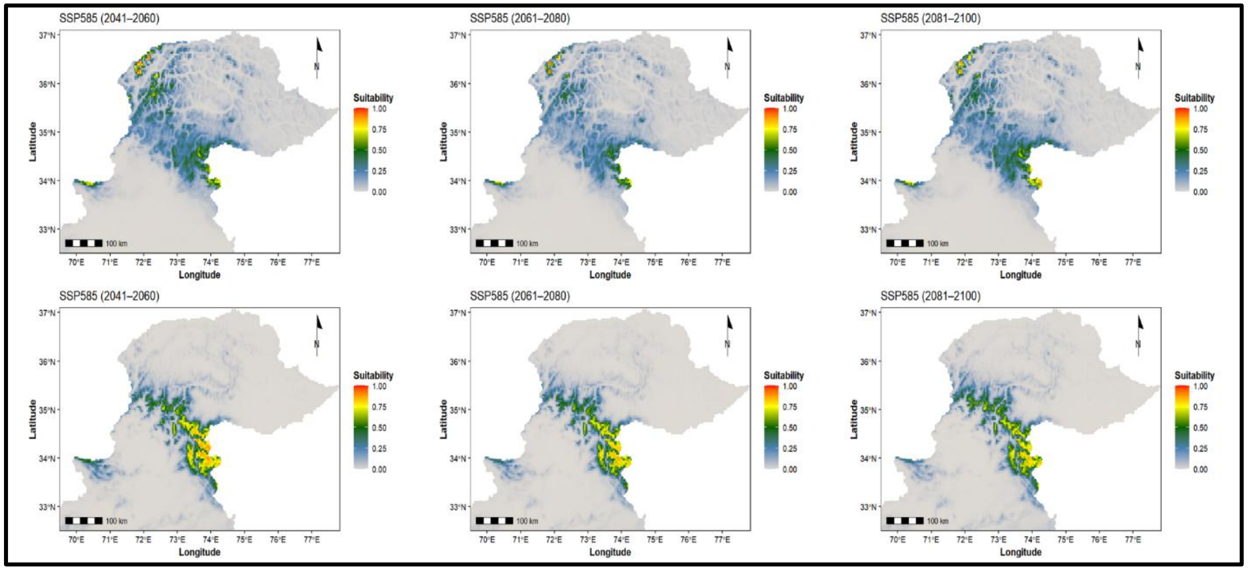


**Figure S4. Habitat suitability under SSP585 shows that both species persist mainly in limited, environmentally buffered areas.** (A-C) Projected suitability for *Gentiana cachemirica* shows strong confinement to isolated high-elevation regions by the end of the century, while (D-F) projected suitability for *G. kurroo* indicates widespread reduction in suitable habitat and increased fragmentation across all periods. Habitat suitability values (0-1) are displayed using a continuous color gradient from low (blue) to high (red), derived from projected climatic conditions. These projections indicate that under strong warming, only areas with relatively favourable local conditions remain suitable, while most of the landscape becomes unsuitable.
